# Tatajuba ― Exploring the distribution of homopolymer tracts

**DOI:** 10.1101/2021.06.02.446710

**Authors:** Leonardo de Oliveira Martins, Samuel Bloomfield, Emily Stoakes, Andrew Grant, Andrew J. Page, Alison E. Mather

**Author notes:** Corresponding author: Leonardo de Oliveira Martins, Corresponding.

## Abstract

Length variation of homopolymeric tracts, which induces phase variation, is known to regulate gene expression leading to phenotypic variation in a wide range of bacterial species. There is no specialised bioinformatics software which can, at scale, exhaustively explore and describe these features from sequencing data. Identifying these is non-trivial as sequencing and bioinformatics methods are prone to introducing artefacts when presented with homopolymeric tracts due to the decreased base diversity. We present tatajuba, which can automatically identify potential homopolymeric tracts and their putative phenotypic impact, allowing for rapid investigation. We use it to detect all tracts in two separate datasets, one of *Campylobacter jejuni* and one of three *Bordetella* species, and to highlight those tracts that are polymorphic across samples. With this we confirm homopolymer tract variation with phenotypic impact found in previous studies and additionally find many more with potential variability. The software is written in C and is available under the open source license GNU GPL version 3 from https://github.com/quadram-institute-bioscience/tatajuba.

## Introduction

The presence of repetitive DNA bases across bacterial genomes is ubiquitous and is associated with important phenotypic changes, especially in organisms with skewed GC content [1,2]. These repetitive regions are known as homopolymeric. Since frameshifts are facilitated by such homopolymeric tracts (HT), they can lead to phase variation; the resultant change can lead to truncation of coding sequences with consequent changes in gene expression and therefore phenotype. Or if the HT occurs in non-coding control regions, it can affect the expression of genes. Identifying and monitoring all HTs in a sample can be challenging due to the large numbers and difficulty in identifying such tracts from sequence data. Since these frameshift events can be common in a population and regulate, or affect, the expression of genes with important phenotypic traits, effort should be directed to identify variation in tract lengths.

To date, evolutionary analyses have focused on single nucleotide polymorphisms (SNPs) and insertions and deletions (indels). In contrast to SNPs, HTs are harder to sequence, depend on the GC-content and may lead to biased coverage by current sequencing technologies [3–6]. Furthermore, they are problematic to align across samples, since HTs are represented as indels (insertion-deletions) in phylogenetics. With the exception of a few specialist models [7–10], indels are treated as missing data in evolutionary inferences [11–13]. They are furthermore challenging to catalogue, unless there is a specific region of interest which can then be curated manually for the presence of HTs. Our proposed algorithm, implemented in the software tatajuba, can be applied to any genome with an available annotated reference sequence, and can extract all HTs within the genome while allowing for tract length polymorphism even within a sample. To account for sequencing errors, tatajuba conservatively only calls HTs in areas with high read coverage, supported by both strands, and with sufficiently long flanking DNA on each read. The flanking regions must allow for the HT to be uniquely mapped to the reference genome, with length defined by the user, currently limited to less than 32bp on each side. We consider read coverage in both forward and reverse strands by using a canonical representation of the tracts. The software can optionally be configured at runtime to be less conservative, with the risk of an elevated false positive rate of HTs identified or losing the ability to map against the reference in a few cases.

Our objective is to fully describe the distribution of tract lengths for all HTs in a sample, the potential phenotype impact of such changes, and compare differences in HT length across a given set of samples. Differences in the variability of HTs across samples can give us information about evolutionary processes (diversity) and phenotype, and can be explicitly modelled. Using the whole distribution across reads, as opposed to assuming a single consensus sequence per sample, allows us to account for minor variants and intra-sample diversity, essential for observing small-scale evolutionary trends and phase variation in clonal populations [14].

We demonstrate the software capabilities on two data sets where the importance of phase variation has been previously described, *Campylobacter* [2] and *Bordetella* [15], but this tool can be applied to any microbial species.

## Materials and methods

The software focuses on describing the tract length distributions across samples, by mapping them to a reference annotated genome and finding those with variability across samples. Given a FASTQ file as input, a HT is found from sequencing reads of nucleotides, and is defined as the same base repeated (*e.g*., 3 times or more), and flanked by a pair of k-mers, called “contexts”. Each context is thus a short DNA segment, between 10 and 32 bps typically, which flanks an HT and contains a rich set of nucleotide sequences. Using these flanking regions to anchor the HT avoids bioinformatic artefacts usually inherent around HTs. We will refer to a HT together with its pair of contexts simply as a tract, and we will consider only those tracts that can be mapped to the reference genome ―otherwise they are discarded, but can be reported to the user since a large fraction of those can be indicative of an inappropriate choice of reference sequence or sequence contamination.

Furthermore, in the presence of paralogs or, specifically, when the exact same tract is mapped to more than one region in the reference genome, the method chooses the first one in genomic coordinates. To avoid inclusion of sequencing errors, we only consider an exact DNA sequence (*i.e*., read segment with identical contexts and homopolymer length) observed in more than a number of reads fixed by the user (default is 3). Afterwards we can merge these identical DNA segments into a tract if their contexts are similar enough (based on their edit distance) and map to the same location in the reference genome, and their homopolymers are composed of the same base (see table 1). Specifically, we use the C functions from BWA-aln for single ended reads [16] to map each tract against the reference. Some tracts will not be mapped due to inappropriate reference or contamination, and these are excluded from further analysis. In both cases the tract is not represented in the reference genome of choice (but may map to a different reference). The tracts might also fail to map due to poor aligner performance on low complexity regions, but large flanking regions should minimise this risk.

**Table 1:**
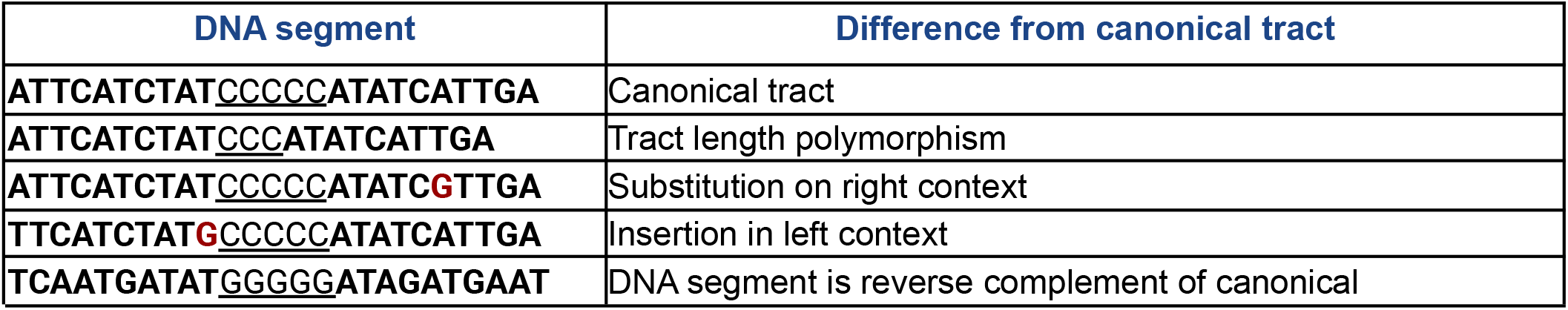
the tract can be variable. All the following read segments come from same tract, represented at the top as the “canonical” or exemplary tract (notice that the contexts have fixed size of 10 bases in this example):

The tracts are therefore comparable across samples, where we can now create a list of tracts present in at least one sample (and also in the reference). In addition to reporting all identified tracts, we also highlight those that present variability across samples or in relation to the reference genome. This will generate a smaller set of tracts with potentially important biological implications. The measure of dispersion used here to find variable tracts is the absolute range (MAX-MIN). Other measures could be used, for instance the *relative difference of ranges* (similar to the coefficient of range), but it is being used here solely to exclude the tracts that do not change at all between samples.

Besides the read files for all samples to be analysed, tatajuba requires the reference genome both in fasta format and its GFF3 file, such that it can access the annotations harbouring each tract and identify if the tract falls within a coding sequence or not. It works with prokka’s GFF3 output [17], which means that the GFF3 file (1) can contain the fasta sequences, and (2) can have more than one contig/chromosome/genome. A multiple sequence FASTA file can also be provided in addition to the GFF3 file, which renders tatajuba compatible with multiple reference genomes.

### Software Implementation

The Levenshtein distance is used to decide if contexts from read segments represent the same tract. There are a few cases where the program detects two tracts that map to the exact same location in the reference genome. These cases may reflect a substitution within the homopolymeric region, which renders the flanking regions too dissimilar to be merged by the program. It can also happen when the flanking region includes a HT itself. Two examples are given in Table 2, where the same sample can present both versions of the tract.

**Table 2:**
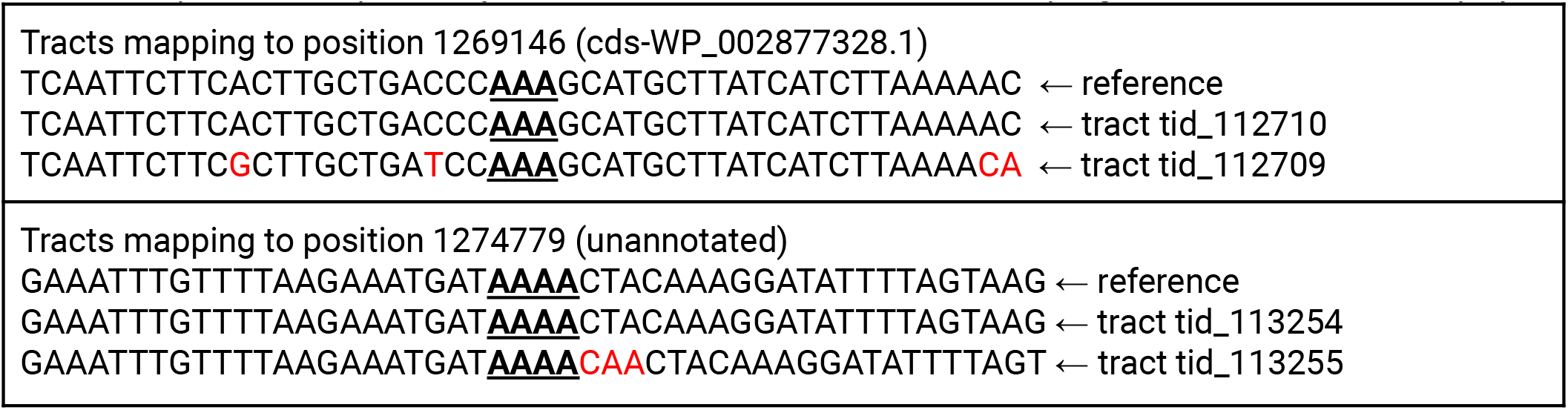
examples where tatajuba finds more than one tract mapping to the same location in the reference genome. The polymeric tract is represented in bold, and differences in context (flanking regions) are highlighted in red. The top panel shows an example where substitutions on the flanking regions are responsible for the classification, while the bottom panel shows potentially successive insertions, with the “C” disrupting the otherwise increased poly-A.

The maximum distance allowed can be controlled by the user, with the default being one, but the example above highlights how this information might be useful. By using the distribution of lengths in contrast to their consensus value, we can observe subtle changes in the populations, as for instance a sample where for a particular tract most reads have a length 4 homopolymer, but a few have length 5.

### Samples

It has been shown that the HT variation can be used to identify particular phenotypic traits, such as cell shape in *Campylobacter jejuni* [2]. We thus analysed 100 *Campylobacter* samples used in [2]: 68 of which had a phase variation described in one of the two genes of interest (*pgp1* and *pgp2*), and 32 “wild type” samples, *i.e*., without a phase variation described in the original paper (list of samples and accession numbers available as Supplementary Material and at https://github.com/quadram-institute-bioscience/tatajuba). We used *C. jejuni* M1 (ASM14870v1) as the reference genome.

Another study described a set of HTs with potential biological relevance and variability across three *Bordetella* species [15]. In this study there were no available data to evaluate intra-species variation, but recently many data sets have been deposited in public repositories. We therefore were able to analyse 108 *Bordetella* samples downloaded from ENA (91 *Bordetella. pertussis*, 7 *Bordetella parapertussis*, and 10 *Bordetella bronchiseptica*), using *B. pertussis* Tohama I (ASM19571v1) as the reference genome. The *B. pertussis* samples come from bioprojects PRJNA348407, PRJNA356412, PRJEB42353, and PRJEB38438 [18–20], while the *B. parapertussis* and *B. bronchiseptica* come from bioproject PRJNA287884 [21].

## Results

From each sample, we selected all tracts with homopolymeric lengths of 3bp or more, which were present in at least 8 reads. We assumed a context of 28 bp on each side,and merged those with a Levenshtein distance smaller than 2. For these parameters, we found a total of 182,983 tracts for *Bordetella* and 144,371 tracts for the *Campylobacter* data set which could be mapped to their reference genomes. The numbers for each individual sample are shown in Figure 1, where we can see that (1) both data sets have comparable numbers of mapped HTs: around 140k tracts for most samples for both data sets, although the variation is higher for *Bordetella*; and (2) a high variation in the number of unmapped tracts, with large values found for *Bordetella*. The unmapped tracts are a combination of the underlying evolutionary processes and artefacts like technical contamination or poor choice of the reference genome. When the genome sizes are taken into account (1.6Mb for *Campylobacter* against 4Mb of *Bordetella*) then the frequency of HTs in *Campylobacter* is almost twice as high as in *Bordetella*.

**Figure 1:**
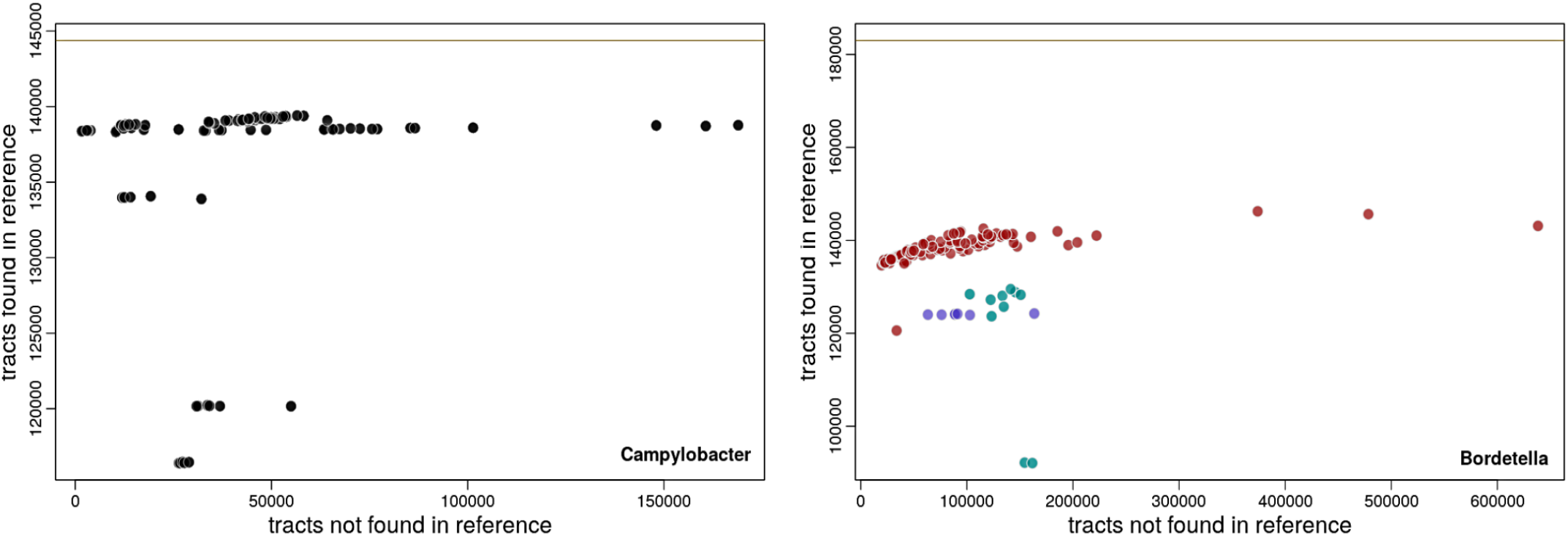
Number of tracts that could be mapped or not to the reference genomes per sample for the *Campylobacter* (right) and *Bordetella* (left) datasets. Each point represents a sample, where some HTs can be mapped back to the reference genome (y axis) and some HTs cannot be mapped (x axis). The total number of HTs found by tatajuba are the sum of the x and y axes for each sample. The beige horizontal lines are the total number of mapped tracts, and represent the union over individual samples. For the *Bordetella* dataset (right panel), the colours represent samples from different species (red for *B. pertussis*, green for *B. bronchiseptica*, and purple for *B. parapertussis*).

Tatajuba analysed the 100 *Campylobacter* samples in 12 minutes and the 108 *Bordetella* in 15 minutes, using a computer with 48 cores and using less than 30GB. Currently we exclude tracts that cannot be mapped to the reference. In the *Campylobacter* dataset, we found 42,832 tracts with variable length distributions, of which 38,295 were annotated, that is, belonged to a gene or RNA. From the *Bordetella* data set we found 129,180 variable tracts, 111,512 of which were in annotated regions. By using the average length of a HT as a feature, we can cluster all samples based on how similar their sets of tracts are (in terms of the average length). The results are shown in Figure 2, where we see that such clustering is in agreement with their evolutionary relationships.

**Figure 2:**
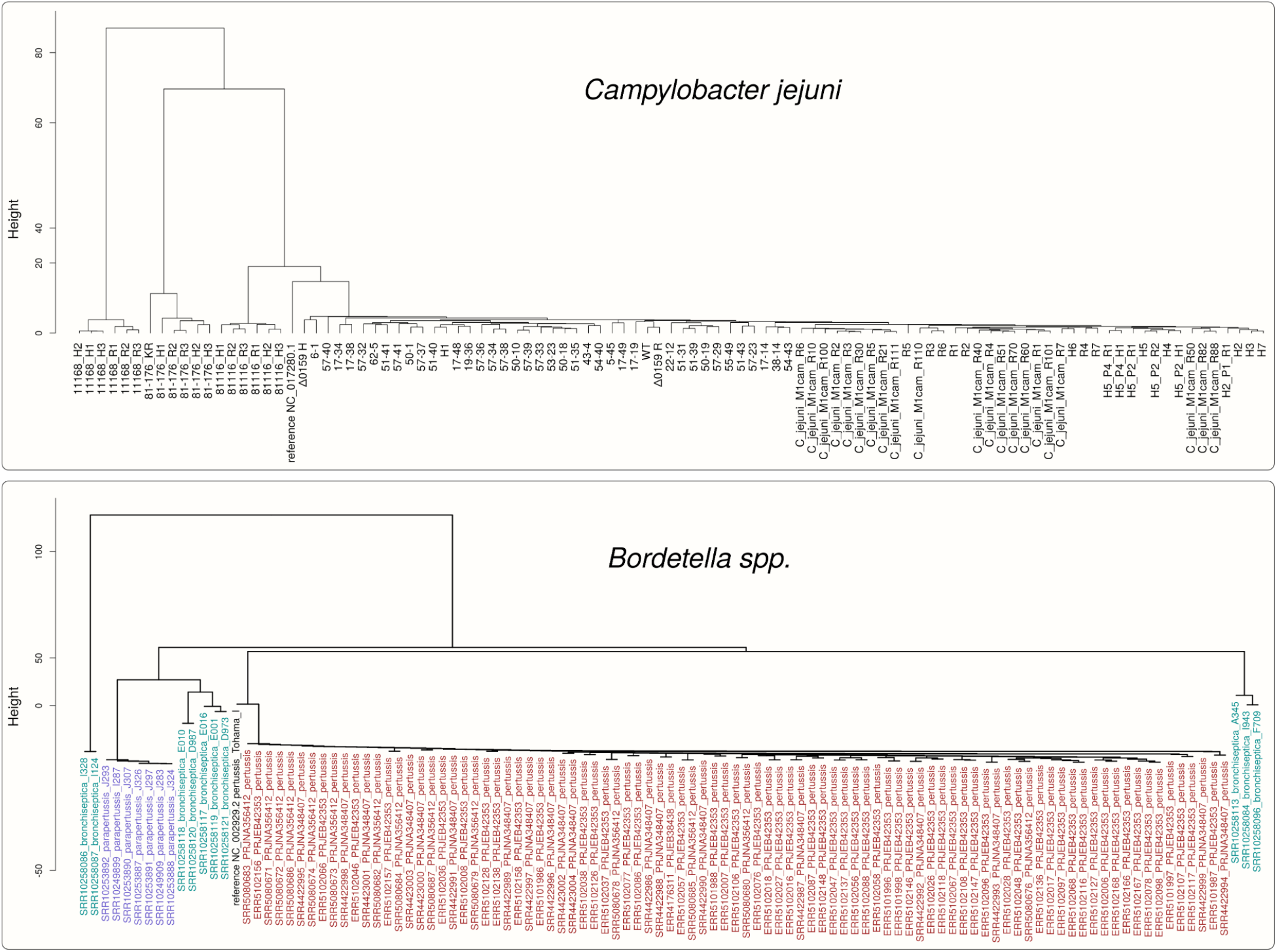
Dendrogram based on tract length profile similarity, using the average tract length as a feature. At the top we have the dendrogram for the *Campylobacter* samples. The bottom dendrogram shows the *Bordetella* samples, with colours representing the species as in Figure 1, with the reference genome (*B. pertussis*) in black.

We furthermore measured the variability of each tract across samples, and visualised them along the genome (Figure 3). In this figure the variability is represented as the maximum difference between tract lengths across samples, where we can see it is not unusual to observe tracts where samples have a length difference higher than 4, for instance. This dispersion measure is currently used only to exclude HTs with no variation at all across samples, but as we see it gives an overview of highly variable tracts. Other measures can be implemented, which are more robust to outliers or are based on the length distribution within a sample instead of a point estimate. Furthermore length changes in multiples of three (a codon) might have a lower impact than those potentially changing the reading frame.

**Figure 3:**
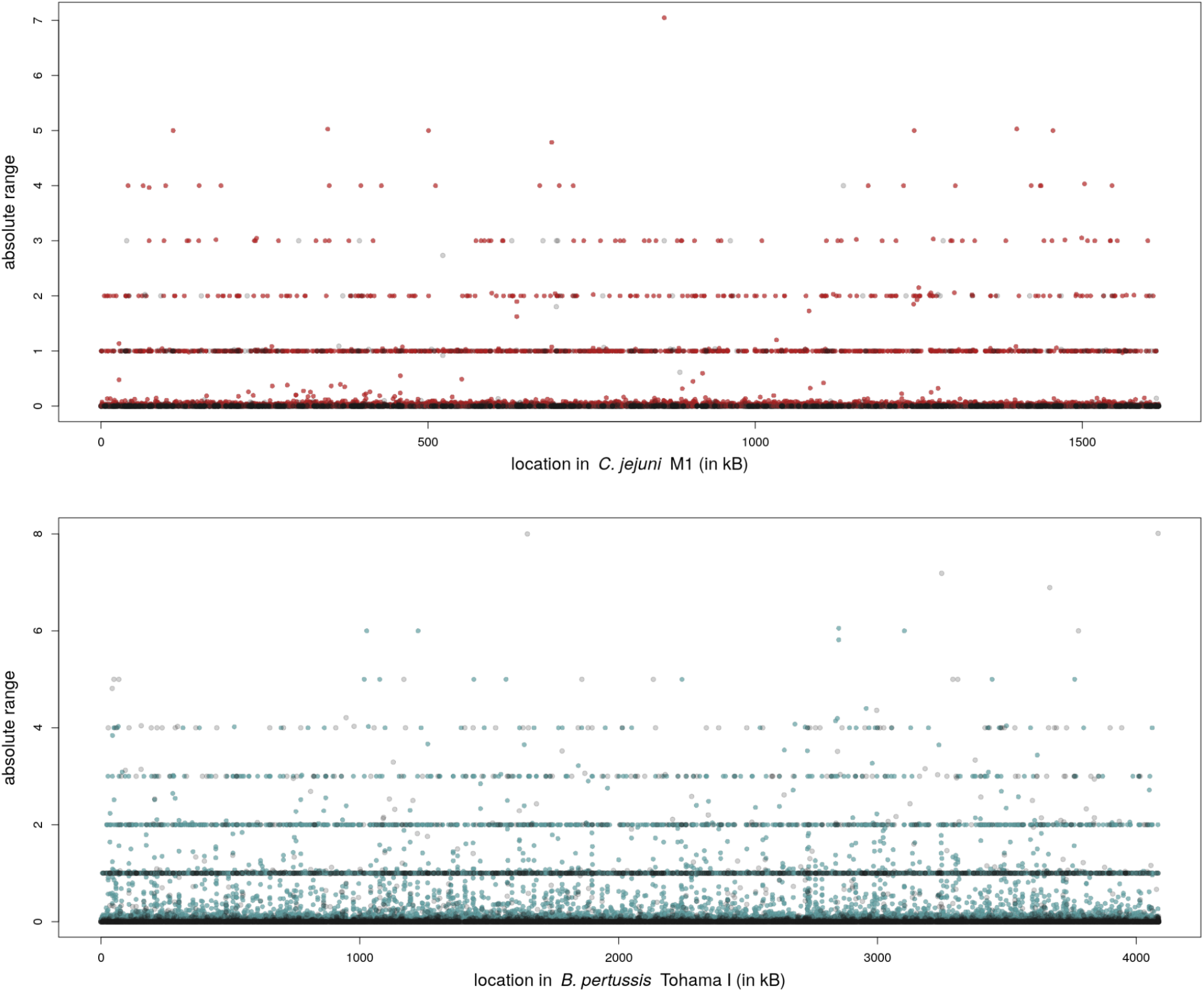
Tract length ranges (maximum minus minimum values across samples) across the reference genome of the two example data sets. Blue or red dots represent tracts in an annotated region while gray dots are not annotated. The tract length is estimated through the average length over reads, and only variable tracts are shown (i.e. those with range higher than zero).

In [2], 18 modifications in genes *pgp1* and *pgp2* were described which were associated with rod-shaped *C. jejuni* (Table 1 of that paper). Of these, 13 involve a change in the length of the homopolymer tract. In Figure 4 we observe that most mutations previously described leading to tract length modifications [2] are also found by tatajuba. The correspondence between the changes found in [2] and in tatajuba are shown in Table 3. Some are missing since we limited the search to homopolymers of length 3 or higher. Interestingly, tatajuba misses the changes originally reported in locations 1268739 and 1268827 from 3A to 2A. Instead, it reports 3A for all samples, since it stores the distribution of homopolymers truncated at 3: even if the 2A (dimer) form is more frequent than the trimer 3A (and would therefore be the consensus), tatajuba does not keep track of dimers. Upon further inspection (by allowing dimers, results not shown) we confirmed this is the case for 1268739. For 1268827 and other cases where tatajuba does not detect the HT change, like locations 1268899 and 1268944, may be due to low tract coverage or missing context from raw reads (see also Figure 6 below for an example where there is insufficient context around a given HT).

**Figure 4:**
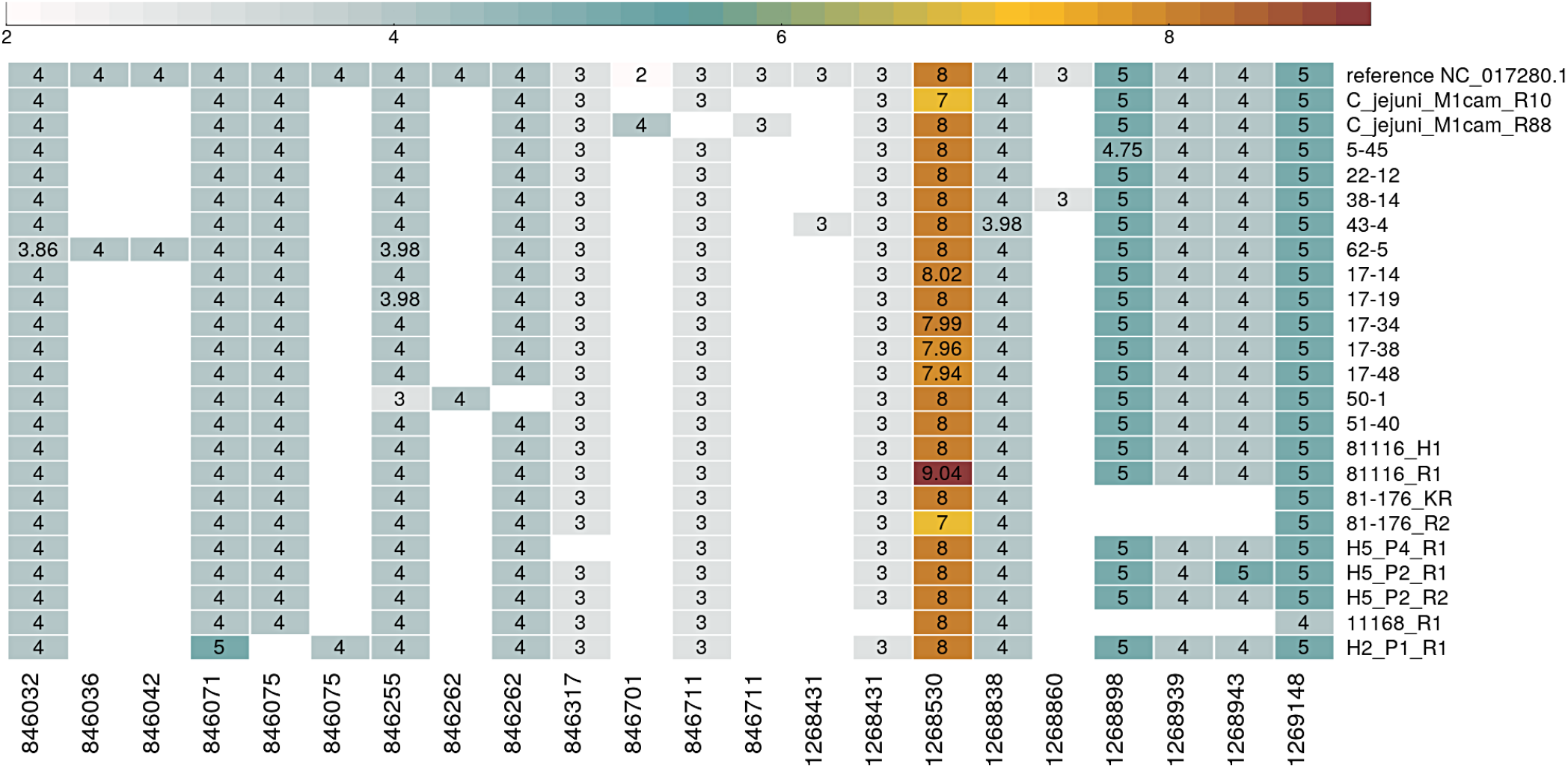
Average tract length for selected *Campylobacter* samples, over genes *pgp1* (from 1268323 to 1269717) and *pgp2* (846020 to 846997) for tracts described in [2]. Rows correspond to samples, and columns are the genomic location of the HTs - only variable tracts are shown i.e. if a tract has same length over all samples, then its column is excluded. The same location appears more than once for cases where tatajuba decides that the contexts are too distinct even if mapping to the same location in the reference genome. Tract lengths smaller than three were not considered by tatajuba and therefore are absent (empty cells).

**Table 3:** correspondence between HT length modification found here and in [2]. The location shown corresponds to the beginning of the homopolymer, with tatajuba starting at zero (instead of one). Tract lengths smaller than 3 were not recorded and thus appear as ‘absent’.

| Location in [2] | Location in tatajuba | change | Samples in Figure 4 where change is observed | Confirmed by tatajuba | Observations |
| --- | --- | --- | --- | --- | --- |
| 846037 (pgp2) | 846032 | 4 T > 3 T | 62-5 | Y |  |
| 846075 (pgp2) | 846075 | 4 A > 3 A | H2_P1_R1 | Y |  |
| 846256 (pgp2) | 846255 | 4 A > 3 A | 50-1 | Y | 62-5 and 17-19 show variability |
| 846319 (pgp2) | 846317 | 3 A > 2 A | H5_P4_R1 | Y |  |
| 846702 (pgp2) | 846701 | 2 G > 4 G | C_jejuni_M1cam_R88 | Y |  |
| 1268531 (pgp1) | 1268530 | 8 A > 7 A | C_jejuni_M1cam_R10, 81-176_R2 | Y |  |
| 1268531 (pgp1) | 1268530 | 8 A > 9 A | 81116_R1 | Y |  |
| 1268739 (pgp1) | 1268738 | 3 A > 2 A |  | N | All samples have length of 3 in tatajuba |
| 1268827 (pgp1) | 1268826 | 3 A > 2 A |  | N | All samples have length of 3 in tatajuba |
| 1268899 (pgp1) | 1268898 | 5 T > 4 T | 5-45 | Y/N | Not found in previously reported 17-48; 5-45 is only partial |
| 1268944 (pgp1) | 1268943 | 4 A > 5 A | H5_P2_R1 | Y/N | Tract is absent in some samples |
| 1269149<br>(pgp1) | 1269148 | 5 A > 4 A | 11168_R1 | Y |  |

A previous study described 58 HTs in *B. pertussis* putatively involved in phase variation [15]. This study employed a Markov model to find HTs longer than expected by chance, using three reference genomes: *B. pertussis*, *B. parapertussis*, and *B. bronchiseptica*. We thus compared the previously identified genome locations with the closest equivalents as reported by tatajuba on our *Bordetella* dataset. The result is shown in Figure 5, where all originally reported HTs are found by tatajuba, with the exception of one at ORF BP0146. In [15] they consider the gene strand when annotating the start of the HT, but to make locations comparable, we report the HT’s location with respect to the leftmost base in the reference.

**Figure 5:**
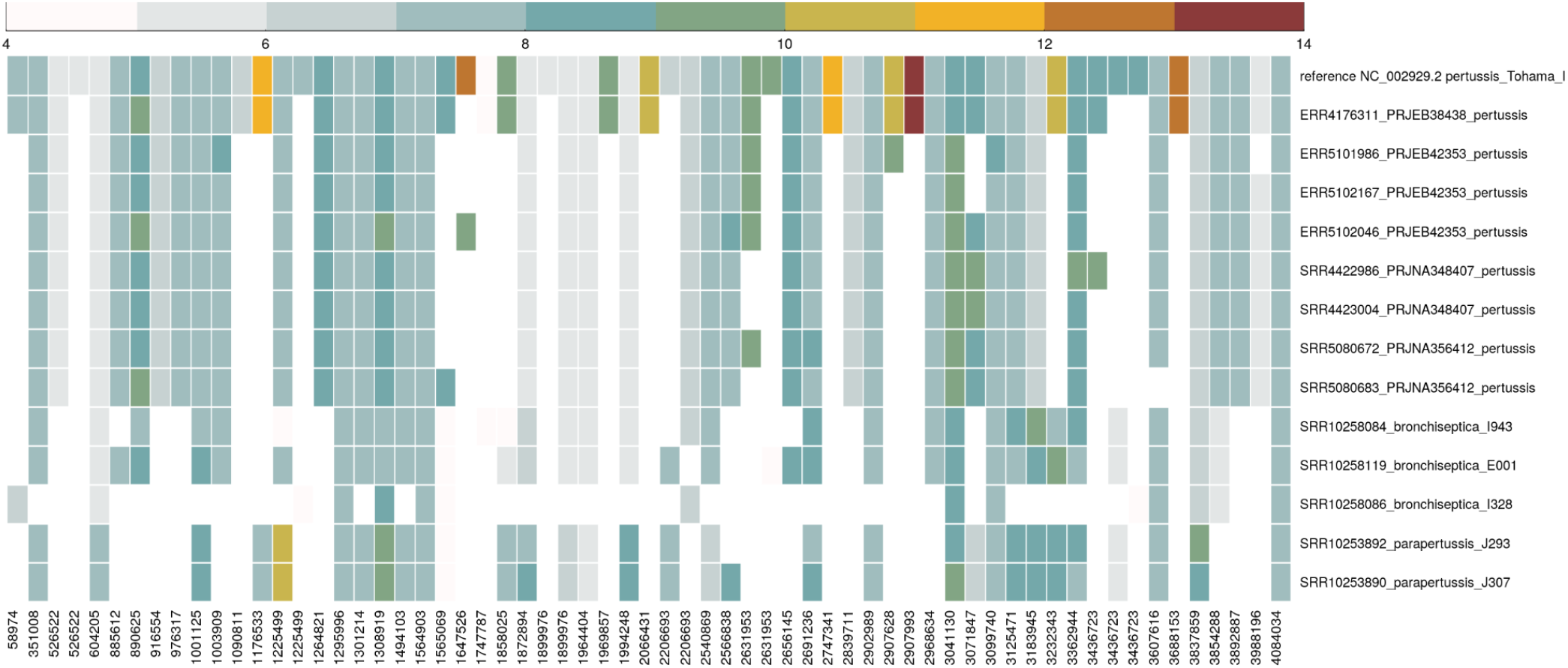
Average tract length for selected *Bordetella* samples, over regions reported in [15]. The samples were arbitrarily selected for display purposes, to show the variety of tract lengths.

## Discussion

One future direction is to use the HTs explicitly as phylogenetic markers, for instance by extending an alignment from their flanking regions. The HTs can be rapidly identified across samples, and as we observed carry evolutionary information. By exhaustively exploring populations of genomes for their presence, we may find regions of phylogenetic importance. The software allows for multiple genome annotation and therefore can work with several reference genomes.

Tatajuba can be used to help infer the phenotypic effect of tract length variations, by finding those in coding regions and by describing the change in tract length ― in coding regions, we expect frameshift mutations to have a higher impact than a tract length difference of multiples of three.

Strand bias, where reads from the forward strand disagree with reads from the reverse strand, are more common around homopolymeric regions [22–24]. Although less affected than other technologies, Illumina sequencing can generate spurious indels within HTs [25–27], especially for HT lengths longer than 14bp [28,29], and it was estimated to affect up to 1% of the genes in a metagenomic data set [30]. A recent survey showed that up to 5.3% of all Illumina errors are related to homopolymers of length 3 or more [31]. To correct for the length errors induced by strand bias, tracts present in only one strand can be flagged [32], or a filter can be added to exclude length disparities associated with the strand [22]. Tatajuba only considers tract lengths observed on both strands. It has been observed that error correction algorithms might introduce errors around HTs [5], although alternatives exist, in particular quality-score-based error removal [5,33].

BWA-aln is optimised for short reads and has very good performance when the read is similar to the reference genome except for HTs at the end of the alignment [34]. However, it fails to map reads with lower matches, unlike mappers which can spuriously return random mappings [34]. BWA-aln is thus well suited for tatajuba: in our case the HT is never at the end of the alignment, since it is flanked by conserved sequences, and we assume the presence of a close reference genome.

It is important to keep in mind that our procedure is based on raw reads (samples), and reports HTs which were observed in these samples. This explains why tatajuba could not find the HT reported at BP0146 (genome location 174886). In Figure 6 we show a list of all reads from five samples potentially containing the HT, as defined by a small stretch of the poly-G followed by a short flanking region of 6 bases. There we see that this HT is likely to be rejected by tatajuba since it does not have sufficient context (*i.e*., flanking regions) or its coverage depth is too low.

**Figure 6:**
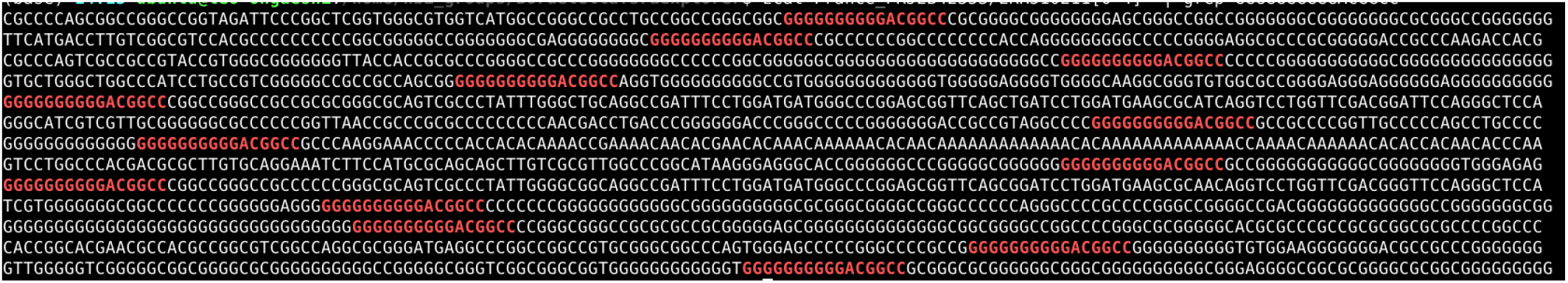
Search for a specific homopolymer tract using raw reads. The HT reported in [15] has 15 Gs in location 147886 of the *B. pertussis* Tohama I genome. A loose text search for sequence GGGGGGGGGGACGGCC (and its reverse complement GGCCGTCCCCCCCCCC) returns the reads above for five samples (ten paired end files), with the search text in red. Each sample read has fewer than 5 valid tracts.

It is also important to note that tatajuba compares the HTs from each sample to a common reference genome and thus the comparison between samples from different species is automatic, that is, we don’t need to map between reference genomes as in [15].

## Conclusion

HTs are widespread in many bacterial species, and variation in HT length can regulate gene expression. In both bacterial species examined here, HTs were found in large numbers, rendering the task of merely identifying such tracts unmanageable without automation. Clearly, with this number of HTs, an automatic/systematic way of investigating variation is required. Although our analyses relied on a single reference genome, we show how we can have meaningful results even when several species are analysed together. Tatajuba provides a huge scope for identifying potential genetic and therefore phenotypic variation which has thus far not yet been explored systematically. It therefore facilitates the discovery of important biological insights. Tatajuba cannot solve the coverage bias induced by some sequencing technologies towards HTs, but it excludes tracts with low depth and within reads without enough context, for instance those at the end of the read, without a flanking region.

Some sequencing platforms are sensitive to homopolymers, which can induce indel errors. For instance, MiSeq can find the correct HT length more often than the Ion Torrent PGM or the 454 GS Junior [25]. When the HT leads to sequencing mistakes, reads from the forward strand may produce a HT length distinct from reads from the reverse strand ―which would be summarised by tatajuba through the length distribution. We cannot fully eliminate systematic bias from sequencing and bioinformatics, but we can limit it. Additionally, tatajuba can be used as a quality control tool to identify these systematic bias issues. It should be used whenever any sequencing method, technology, or library preparation is being updated on a standard set of bacteria.

Tatajuba is available under the open source GNU GPL 3 licence from https://github.com/quadram-institute-bioscience/tatajuba. The software is written in ANSI C (C11 standard with GNU extensions), validated using unit tests and packaged for autotools and Conda.

## Acknowledgements

The authors gratefully acknowledge the support of the Biotechnology and Biological Sciences Research Council (BBSRC); this research was funded by the BBSRC Institute Strategic Programme Microbes in the Food Chain BB/R012504/1 and its constituent projects BBS/E/F/000PR10348 (Theme 1, Epidemiology and Evolution of Pathogens in the Food Chain) and BBS/E/F/000PR10349 (Theme 2, Microbial Survival in the Food Chain) and BBS/E/F/000PR10352 (Theme 4, Research Infrastructure), and by the Quadram Institute Bioscience BBSRC funded Core Capability Grant (project number BB/CCG1860/1).

